# Phage Display Broadly Identifies Inhibitor-reactive Regions in von Willebrand Factor

**DOI:** 10.1101/2021.03.08.434464

**Authors:** Andrew Yee, Manhong Dai, Stacy E. Croteau, Jordan A. Shavit, Steven W. Pipe, David Siemieniak, Fan Meng, David Ginsburg

**Author notes:** Corresponding authors:David Ginsburg, University of Michigan, Life Sciences Institute, 210 Washtenaw Ave Ann Arbor, MI 48109, Andrew Yee, Baylor College of Medicine, Feigin Tower, 1102 Bates St. Rm. C.1025.09, Houston, TX 77030.

## Abstract

**Background:** Correction of von Willebrand factor (VWF) deficiency with replacement products containing VWF can lead to the development of anti-VWF alloantibodies (i.e., VWF inhibitors) in patients with severe von Willebrand disease (VWD).

**Objective:** Locate inhibitor-reactive regions within VWF using phage display.

**Methods:** We screened a phage library displaying random, overlapping fragments covering the full length VWF protein sequence for binding to a commercial anti-VWF antibody or to immunoglobulins from three type 3 VWD patients who developed VWF inhibitors in response to treatment with plasma-derived VWF. Immunoreactive phage clones were identified and quantified by next generation DNA sequencing (NGS).

**Results:** NGS markedly increased the number of phage analyzed for locating immunoreactive regions within VWF following a single round of selection and identified regions not recognized in previous reports using standard phage display methods. Extending this approach to characterize VWF inhibitors from three type 3 VWD patients (including two siblings homozygous for the same *VWF* gene deletion) revealed patterns of immunoreactivity distinct from the commercial antibody and between unrelated patients, though with notable areas of overlap. Alloantibody reactivity against the VWF propeptide is consistent with incomplete removal of the propeptide from plasma-derived VWF replacement products.

**Conclusion:** These results demonstrate the utility of phage display and NGS to characterize diverse anti-VWF antibody reactivities.

## Introduction

Plasma von Willebrand factor (VWF) stabilizes coagulation factor VIII (FVIII) and directs platelets to sites of vascular injury[1, 2]. Abnormalities of VWF result in several types of von Willebrand disease (VWD)[3]. Intravenous infusion of plasma-derived VWF concentrate is the current standard of care for pediatric VWD patients with significant bleeding and poor response to desmopressin[3, 4]. However, the development of anti-VWF alloantibodies (i.e., VWF inhibitors), which primarily occurs in type 3 VWD patients (prevalence of ~1:1,000,000 of the general population), significantly complicates therapy[5, 6]. In contrast to the high rate of FVIII inhibitor development in severe hemophilia A patients (~30%), alloimmunization to exogenous VWF is less common (~5-10% of type 3 VWD patients) and less well studied[5, 7, 8].

Mature, plasma VWF enters the blood after proteolytic removal of its propeptide and circulates as large multimers, with specific functions localized to distinct domains within the monomer subunits[9]. Previous reports using proteolytic and recombinant fragments of VWF have demonstrated the presence of alloantibodies that recognize the VWF platelet-binding domain and inhibit VWF binding to its platelet receptor, glycoprotein Ib[10–13]. Heterogeneous immunoreactivity to other VWF domains between patients suggests that alloantibody epitopes and potential functional consequences may vary widely among VWD patients with inhibitors[10, 12].

Phage display is a high content method for studying protein interactions. In phage display, bacteriophage are engineered to fuse a coat protein with a library of amino acid variants or random protein fragments of an antigen, thereby linking the expressed peptide with its encoding DNA[14]. Following selection for a specific property (e.g., antibody binding), clones are identified by DNA sequencing. This approach has been used to localize immunoreactive regions in VWF and FVIII for anti-VWF antibodies generated as research reagents and FVIII inhibitors found in patients with hemophilia A, respectively[15–18]. However, the practical limits of standard Sanger DNA sequencing of individual, selected phage impede a comprehensive analysis of an antigenic landscape. Next generation DNA sequencing (NGS) has extended the utility of display technologies (e.g., phage display) for epitope mapping with greater throughput and finer amino acid sequence resolution[19, 20].

We previously reported the application of NGS combined with phage display to provide a comprehensive picture of ADAMTS13 interaction with its target sequence within the VWF A2 domain[21]. We now extend this approach of combining phage display with NGS to identify the segments within VWF recognized by anti-VWF antibodies.

## Materials and Methods

### DNA Constructs

The sequences of all oligonucleotides (P1-P12, IDT DNA Technologies) are listed in Table 1. A FLAG tag (NH_2_-DYKDDDDK-COOH) was inserted into the phagemid pAY-E to facilitate specific elution of bound M13 with enterokinase[22]. Oligonucleotides P1 and P2 were annealed (65.0°C) and ligated into the NotI and SgrAI sites of pAY-E to introduce the FLAG tag. Subsequently, the CcdB-CmR cassette (ThermoFisher) was PCR amplified with primers P3 and P4 and ligated into the SfiI and NotI restriction sites. The resulting phagemid, pAY-FE (GenBank MW464120), was maintained in in TOP10 CcdB Survival T1^R^ cells (ThermoFisher). The sequence of pAY-FE verified by Sanger sequencing.

**Table 1.**
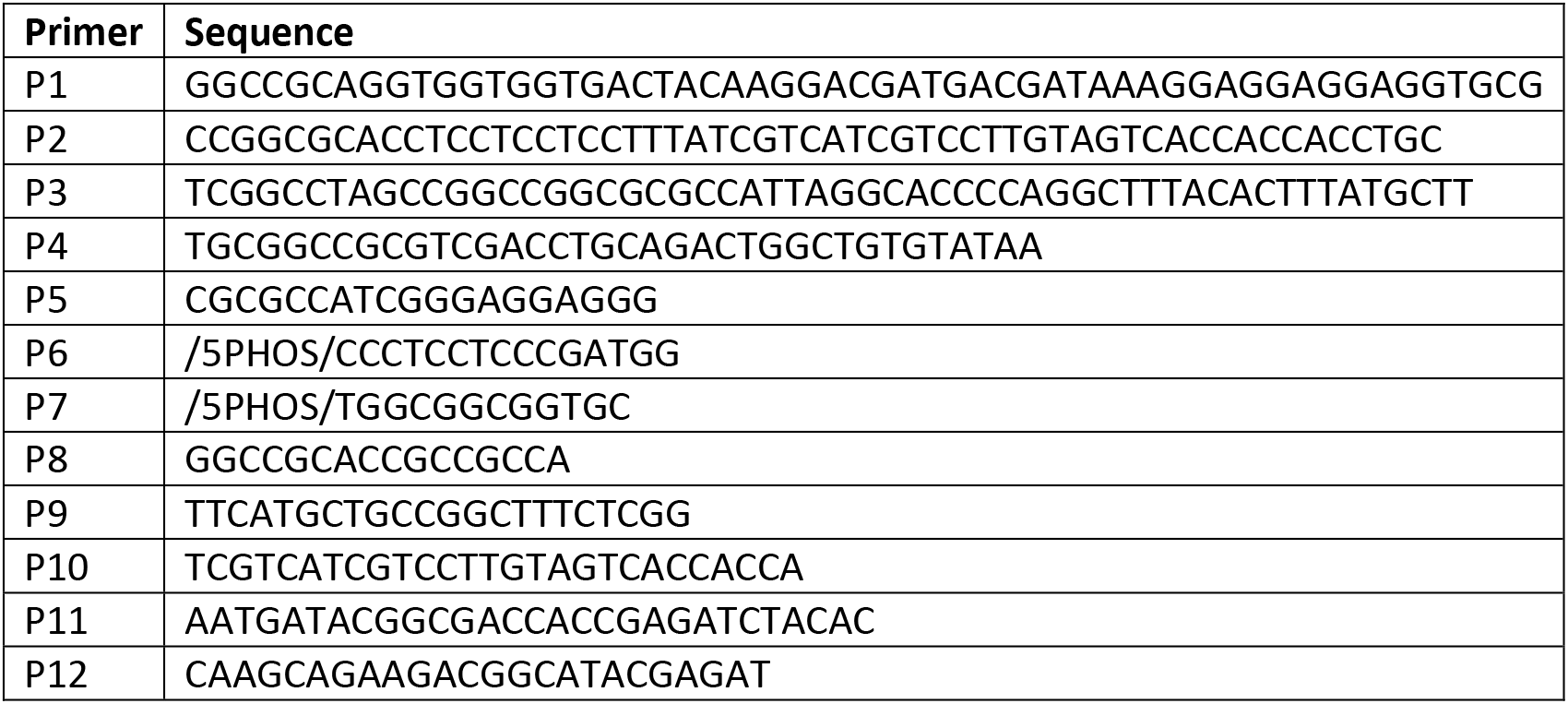
Oligonucleotides.

### Phage Display VWF Fragment Library

The VWF cDNA subcloned into pBluescript was randomly fragmented by sonication. DNA ends were repaired and phosphorylated with the NEBNext End Repair kit (New England Biolabs). Adaptors were created by annealing oligonucleotides (P5/P6 and P7/P8, both at 50.0°C) and were ligated onto the repaired VWF cDNA fragments. Adapted fragments were fractioned across a 2% (w/v) agarose gel, collected (~300bp to ~1,000bp), purified by electroelution and phenol/chloroform extraction, and ligated into pAY-FE at the AscI and NotI sites. Ligation products were transformed into XL1-Blue MRF’ cells (Agilent) by electroporation in 0.1cm cuvettes at 1.8kV, 200Ω, and 25μF (Gene Pulser, Biorad). Transformed cells were recovered in SOC medium (ThermoFisher) at 37°C for 1h and quantified for ampicillin resistance on LB-AG (Luria broth + 100μg/mL ampicillin + 2% glucose) agar; the VWF fragment library depth was ~2.78×10^6^ independent clones. This library (~12.5mL) was expanded by inoculating 300mL LB-AG and culturing at 37°C with aeration overnight. The expanded library was stored in fresh LB-AG + 20% glycerol as 1mL aliquots at −80°C.

Only a subset of the random VWF cDNA plasmid (in Bluescript) fragments cloned into the pAY-FE phagemid vector would be expected to express a VWF peptide fused to the N-terminus of phage coat protein, pIII (Figure S1A). Display of a VWF protein fragment requires that its subcloned, encoding cDNA be composed of VWF sequence (total plasmid content = ~3/4 VWF + ~1/4 pBluescript) whose orientation matched the phagemid orientation (1/2 of fragments) and whose 5’-end was the first nucleotide of a VWF codon (1/3 of fragments) and whose 3’-end was the second nucleotide of a VWF codon (1/3 of fragments). As a result, only ~1/24^th^ (3/4 × 1/2 × 1/3 × 1/3) of the library is expected to display a VWF protein fragment. The remaining 23/24^th^ of clones are expected to encode non-VWF peptides, premature stop codon(s), or frameshift(s) that prevents expression of a C-terminal pIII fusion.

To generate phage virions displaying VWF fragments for screening, 100mL LB-AG was inoculated with a 1mL aliquot of the VWF fragment phage display library and cultured at 37°C with agitation until mid-log phase (OD_600nm_ = 0.3 to 0.5). Cultures were then infected with M13KO7 helper phage (GE Healthcare Life Sciences) at a multiplicity of infection (MOI) ≥ 10 at 37°C with agitation for 1h. Cells were isolated by centrifugation, resuspended in 100mL 2x yeast extract tryptone (2xYT) media containing 100μg/mL ampicillin and 50μg/mL kanamycin, and cultured at 30°C overnight with agitation. Cells were removed from the culture by centrifugation at 4,000g and 4°C, and phage were precipitated from the supernatant with 2mL PEG/NaCl (16.7% PEG 8000 + 19.5% NaCl) per 10mL supernatant at 4°C. Precipitated phage were collected by centrifugation at 20,000g and 4°C, resuspended in TBS-T (50mM Tris-HCl, pH = 7.5, + 150mM NaCl + 0.05% Tween-20), and reprecipitated with PEG/NaCl (2mL per 10mL supernatant) at 4°C. Phage were finally collected by centrifugation at 20,000g at 4°C and resuspended in antibody selection buffer (TBS-T + 5% bovine serum albumin (BSA)). Phage concentrations were quantified by infecting XL1-Blue MRF’ cells as previously described[22].

### Screening Phage Displayed VWF Fragments for Binding Anti-VWF Antibodies

Magnetic protein G beads (New England Biolabs) were blocked in antibody selection buffer and used to immobilize 10-15μg polyclonal rabbit anti-VWF (Dako A0082, lots 00045319 and 00051141) or immunoglobulin G (IgG) in platelet poor plasma (PPP) from patients diagnosed with an inhibitor against VWF. Following informed consent with protocols approved by Institutional Review Boards of the University of Michigan and of Boston Children’s Hospital, PPP containing anti-VWF alloantibodies were collected from previously described pediatric patients diagnosed with type 3 VWD and inhibitors to plasma-derived VWF concentrate[23, 24]. PPP was prepared by centrifugation as previously described[1]. IgG-bound beads were washed with antibody selection buffer and subsequently incubated with 10^11^-10^12^ phage from the VWF fragment library at room temperature for at least 1.5h. Beads were then washed with antibody selection buffer followed by antibody selection elution buffer (5% BSA + 20mM Tris-HCl, pH = 7.4, + 50mM NaCl + 2mM CaCl_2_ + 0.005% Tween-20). Bound phage were eluted with 20ng/mL enterokinase (New England Biolabs) at 4°C overnight. The supernatants containing the eluted phage were collected and stored at 4°C. The number of phage virions eluted was quantified as previously described[22].

### Phage Identification

Single stranded DNA from 10^5^-10^7^ selected and unselected phage was liberated by proteinase K digestion in digestion buffer (100mM Tris-HCl, pH=8.5-8.8, + 5mM EDTA + 200mM NaCl + 0.2% SDS) overnight at 37°C and further purified with phenol/chloroform/isoamyl alcohol extraction[25]. The region containing random fragments of the VWF plasmid was made double stranded using primers P9/P10 (annealing temperature = 65.0°C) and Herculase II (Agilent) with 16-20 rounds of PCR amplification (Figure S1B). The PCR products were purified with AMPure XP beads (Beckman Coulter) and fragmented by 5 cycles of sonication on a Bioruptor (per cycle: power = high, interval = 30s on/30s off, 5min; Diagenode) to facilitate efficient DNA clustering onto the Illumina sequencing platform. Fragmented PCR products were repaired and dA-tailed using the NEBNext End Repair/dA Tailing kit (New England Biolabs) according to the manufacturer’s instructions and subsequently barcoded with Illumina-compatible, barcoded adaptors (NEXTflex DNA Barcodes, Bioo Scientific) using the NEBNext Ultra Ligation kit (New England Biolabs). The adapted PCR products were purified twice with AMPure XP beads and subsequently PCR amplified for 8 cycles with primers P11/P12 (annealing temperature = 60.0°C). The resulting sequencing libraries were quantified with PicoGreen (Thermo Fisher) and analyzed on a Bioanalyzer (Agilent) according to the manufacturers’ instructions. Sequencing was performed on the MiSeq platform (Illumina) using the 2×75bp kit or on the HiSeq 2500 platform (100 base, paired end; Illumina) with the libraries clustered at 12-15pM and mixed with 5% PhiX genomic DNA (Illumina).

### Computational and Statistical Analyses

We used a custom Python script to process the sequencing data. Our script 1) combined paired reads into one sequence, 2) filtered sequences for junctions between the phagemid and the parental VWF plasmid, 3) mapped the adjoined VWF plasmid sequences, and 4) generated a summary of counts for the nucleotide locations within the parental VWF cDNA plasmid that were found immediately adjacent to the phagemid. The summary data were used as input for the subsequent statistical analyses. The bioinformatic, statistical, and graphing scripts are available on GitHub (https://github.com/yeea00/anti-VWF_Phage.git). Each phagemid/VWF nucleotide fragment was considered as an independent variable. Counts for the first and last VWF nucleotide within DNA fragments containing a phagemid/VWF fusion (Figure S1B) ranged from 0 to ~41,000 and were analyzed with DESeq2 to calculate the fold changes (selected vs. unselected), the mean nucleotide counts (between selected and unselected), and the p-values adjusted for multiple comparisons (q-values) by false discovery rate (FDR) correction[26]. A q-value < 0.1 was considered significant.

### Clearance Studies

Patients II-1 and II-2 were challenged with an intravenous infusion of plasma-derived VWF containing FVIII (Wilate, Octapharma) at 50 IU/kg. VWF antigen, VWF ristocetin cofactor activity, and FVIII activity were clinically evaluated at the indicated time points. A similar clearance study of plasma-derived VWF for patient I-1 has been previously reported[24].

## Results and Discussion

Characterization of the unselected library with NGS showed a full representation of VWF N- and C-terminal nucleotides with comparable frequencies (defined as the number of reads for a nucleotide per total number of reads for all nucleotides), demonstrating the absence of a strong selection for or against any particular VWF cDNA fragment in *E. coli* (Figure S2A). Median frequencies were similar between terminal nucleotides of different VWF reading frames, though the distributions of nucleotide frequencies were different among the three frames (Anderson-Darling p < 1×10^−130^ for both replicates of sense nucleotides and p < 0.05 for both replicates of antisense nucleotides, Figure S2B). These nucleotide frequencies were well correlated between biological replicates of unselected phage (Pearson R = 0.85 and 0.93 for nucleotides from the sense and antisense strands of the parental VWF plasmid, respectively; Figure S2C), demonstrating comparability between different preparations of unselected phage.

In our previous analysis of 81 individual phage clones (32 unique) using Sanger sequencing, we identified 8 distinct regions in VWF (E1-E8 in Figure 2) that bind to a commercial anti-VWF antibody[16]. In this report, we expanded our analysis to 10^5^-10^7^ independent phage with NGS and identified a much more extensive immunoreactivity spanning the entire mature VWF sequence for the same commercial antibody. Following a single round of selection, 1,758 N-terminal and 1,466 C-terminal VWF residues were significantly enriched (fold change ≥ 1.5, FDR adjusted p-value < 0.1, Figure 1). In the absence of anti-VWF antibodies, no specific VWF N- or C-terminal residues were strongly enriched (Figure 1).Comparison of fold-enrichment by 2 different lots of the same commercial anti-VWF antibody resulted in a high degree of agreement (Pearson R =0.96 for N-terminal residues, Pearson R = 0.93 for C-terminal residues). The identified antibody-binding fragments localize exclusively to the mature VWF sequence, with no significantly enriched segments of the VWF propeptide (Figure 2), consistent with the use of plasma-derived VWF as the immunogen[27]. Of note, the application of NGS required fragmentation of the ~400bp-~1,100bp phage insert amplicons to ~100bp-~600bp segments (Figure S1B). As a result, the specific 5’ and 3’ termini for any given fragment cannot be linked, limiting the ability to define minimal, contiguous immunoreactive sequences within VWF. Advances in NGS technologies that enable longer reads together with increased sequencing read depth should help circumvent this limitation in future experiments.

**Figure 1.**
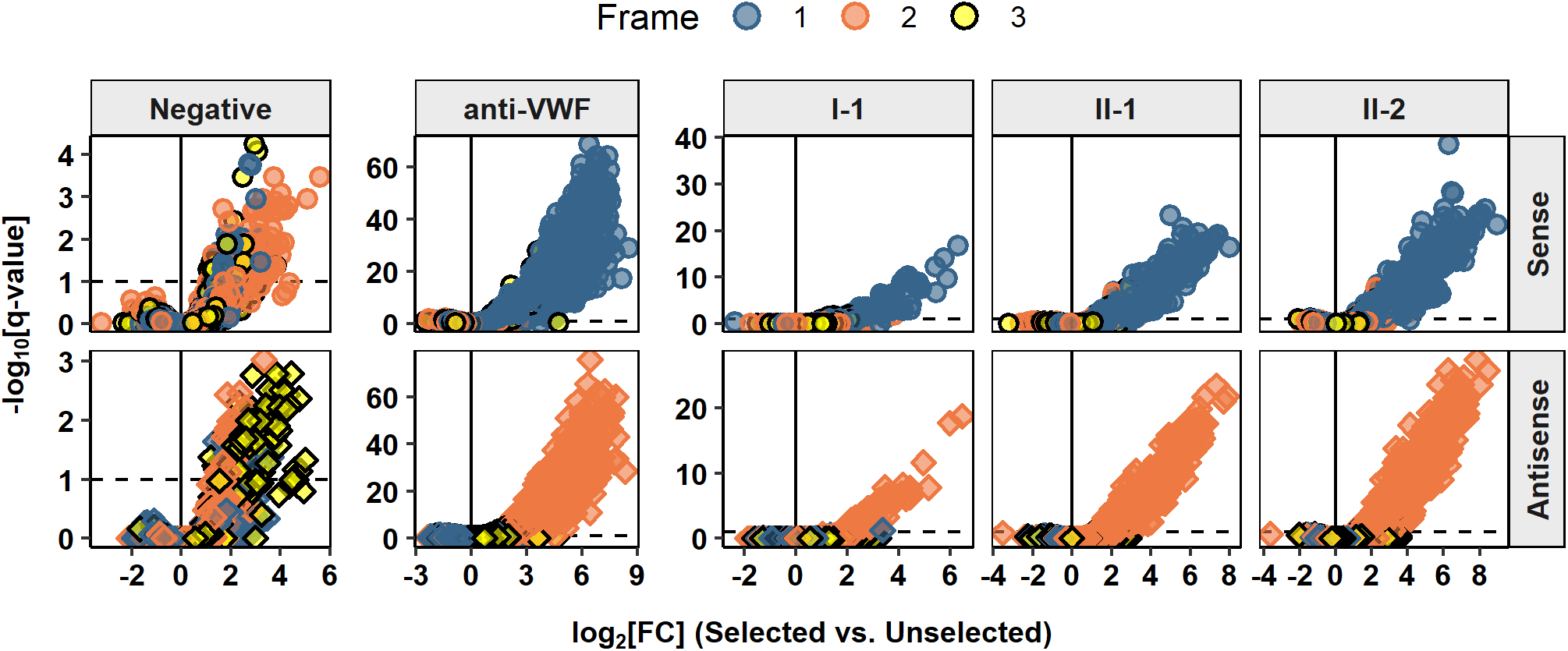
Anti-VWF antibodies significantly enriched N--and C-terminal nucleotides with the appropriate orientation and frame, confirming the phagemid architecture needed to display a VWF protein fragment. Sequences contiguous with the phagemid were identified and quantified for unselected phage and for phage selected in the absence (Negative) or presence of the commercial antibody (anti-VWF) or patient alloantibodies (I-1, II-1, and II-2). Orientation (sense and antisense) and frame (1, 2, or 3) are referenced to the position within the VWF coding sequence. Each marker indicates the VWF nucleotide adjoining the phagemid sequence. Phage displayed VWF peptides are encoded by Sense, Frame 1 and Antisense, Frame 2 nucleotides. Fold-changes (FC, selected vs. unselected) and q-values (p-values adjusted for multiple comparison) were calculated in DESeq2[26]. The dashed line marks the significance threshold (FDR < 0.1).

**Figure 2.**
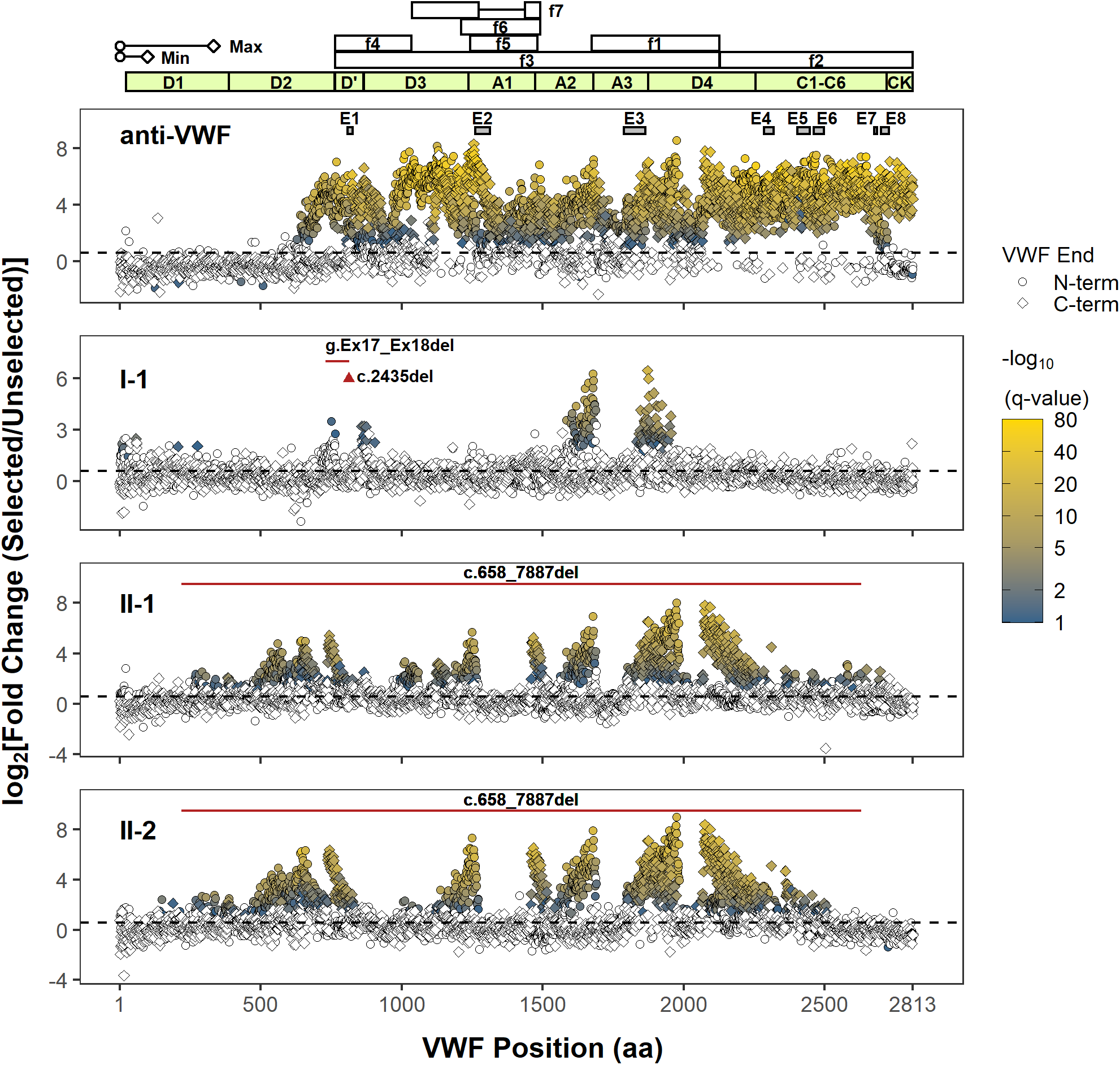
NGS analysis of phage display identifies immunoreactive regions in VWF. Only the position of the first (N-terminal) or last (C-terminal) residue within the identified VWF fragment is shown. The approximate minimal (Min, ~100 amino acids) and maximal (Max, ~333 amino acids) lengths of phage displayed VWF peptides are determined from the size range of VWF cDNA fragments subcloned into the phagemid. However, the phase of N-and C-termini of individual VWF fragments (i.e., which N-terminus is paired with which C-terminus) cannot be determined due to the fragmentation of PCR amplicons of phage DNA for NGS, limiting the delineation of unique epitopes. The coordinates of proteolytic VWF fragments, f1-f7, used to epitope map anti-VWF alloantibodies in prior reports are shown (f1: G1674-E2128; f2: E2129-K2813; f3: S764-E2128; f4: S764-R1035; f5: L1243-G1481; f6: V1212-K1491; f7: K1036-R1274, A1437-K1491)[10–13]. The VWF domain boundaries are annotated in light green. The black dashed line denotes a fold change of 1.5. Terminal residues of phage displayed VWF fragments are marked (VWF End) according to their location and enrichment. Only VWF fragment termini with a significant fold change (FDR adjusted p-value < 0.1) are indicated in shading from blue to yellow, which scales with the significance of fold change (−log_10_(q-value)). A polyclonal anti-VWF antibody (anti-VWF) recognizes fragments spanning the mature VWF sequence. Epitopes, E1-E8, were previously identified by conventional phage display for the same antibody and are shown as gray boxes[16]. VWF inhibitors from type 3 VWD patients (I-1, II-1, and II-2) bind distinct regions in VWF. The location of the patients’ genetic deletion(s) is marked according to the corresponding VWF codon (red triangle for point deletion or red solid line for contiguous deletion). Note that the clustering of significantly enriched termini supports localization of a series of reactive VWF fragments whose internal sequences are graphically depicted as gaps between the clusters (e.g., the immunoreactive region in A3 for I-1 is bounded by clusters of significantly enriched N-and C-terminal residues on the left and right, respectively).

We next applied this approach to localize the VWF binding regions of anti-VWF alloantibodies that developed in three type 3 VWD patients who were treated with VWF replacement products (Figures 1 and 2). Plasma IgG from patient I-1 (compound heterozygous for *VWF* c.2435del and for a deletion in *VWF* of at least 8.7 kb spanning exons 17 and 18[24]) selected VWF fragments that localized to the D’ and A3 domains, with marked enrichment for the A3 domain (Figure 2). Consistent with the lack of selection for VWF fragments spanning the platelet-binding A1 domain, patient I-1’s alloantibodies did not inhibit the ristocetin cofactor activity of infused VWF[24]. Patients II-1 and II-2, brothers both homozygous for a large deletion in *VWF* (c.658_7887del)[23], developed inhibitors with highly similar profiles in the selection of phage-displayed VWF fragments. Similar immunoreactivity to the D3, A1, A3, and D4 domains of mature VWF were found in both patients (Figure 2). Immunoreactivity to the D1 and D2 domains was also identified for both brothers (Figure 2), consistent with the presence of the VWF propeptide in their treatment (plasma-derived VWF derived from several different pharmaceutical sources). In comparison to previous studies that used proteolytic fragments of mature VWF, f1-f7 (Figure 2), to identify regions where anti-VWF alloantibodies bind[10–13], phage display unmasks distinct immunoreactive regions with greater detail for VWF inhibitors from II-1 and II-2. For example, fragment f1 contains the immunoreactive sequences found in A3 and D4 for II-1 and II-2 – note the two gaps between the enriched N- and C-termini under A3 and D4 (Figure 2) – and cannot be expected to distinguish these two distinct regions. The diffuse immunoreactivity observed for II-1 and II-2 suggests the potential for broader inhibition of VWF function than in patient I-1. Consistent with this hypothesis, VWF ristocetin cofactor activity was undetectable in patient II-1 at 30min post-infusion of VWF concentrate despite detectable though low levels of VWF antigen at that time point (Table 2). Rapid clearance of replacement VWF is the likely cause of the failure to restore VWF-dependent hemostasis in these patients; infused VWF was completely cleared within 30min for patients I-1[24] and II-2 (Table 2) and within 3h for patient II-1 (Table 2).

**Table 2.** Clinical Laboratory Evaluation.

| **Patient** | **Test** | **Pre-infusion** | **30min post-infusion** | **3h post-infusion** |
| --- | --- | --- | --- | --- |
| **II-1** | VWF Antigen (%) | 0 | 12 | 0 |
|  | VWF Ristocetin<br>Cofactor Activity (%) | 0 | 0 | 0 |
|  | FVIII (%) | 3 | 59 | 7 |
| **II-2** | VWF Antigen (%) | 0 | 0 | 0 |
|  | VWF Ristocetin<br>Cofactor Activity (%) | 0 | 0 | 0 |
|  | FVIII (%) | 3 | 30 | 5 |

Prior reports that utilized proteolytic VWF fragments to broadly map the immunoreactive regions of VWF inhibitors from a total of 6 unrelated patients concluded that target specificity is unique to each patient[10–13]. We similarly conclude that unrelated patients may have different patterns of immunoreactivity. However, the mechanism(s) for the strikingly similar immunoreactive profiles between patients II-1 and II-2 is unclear. VWF inhibitor development is a complex trait that is unlikely to depend entirely on the VWD-causing variant(s). Not all patients with large genomic deletions in *VWF* develop VWF inhibitors[28]. Furthermore, while patient I-1 and a previously reported patient with the same c.2435del variant in *VWF* both developed anti-VWF alloantibodies, other type 3 VWD patients with this variant (homozygous and heterozygous) do not have VWF inhibitors[28, 29]. Although patients with type 3 VWD have a higher risk for developing VWF inhibitors, alloantibodies have been reported in a patient with type 2B VWD (p.R1308C) who was treated with plasma-derived VWF, demonstrating that a complete lack of VWF expression is not the only risk factor[6].

Notwithstanding the unknown mechanisms that drive anti-VWF alloantibody formation and epitope selection for the patients in this report, we demonstrate a high content approach to broadly survey their alloantibodies’ anti-VWF immunoreactivity. Future refinement of our approach with other molecular techniques may help delineate minimal epitopes in VWF and identify sequences in VWF that may be engineered to be less immunogenic.

## Acknowledgements

We thank the patients for their contributions. NGS was performed by the University of Michigan DNA Sequencing Core. Funding was provided by the National Institutes of Health (HL039693 and HL135793, DG; HL150784 and HL125774, JAS), American Society of Hematology Faculty Scholar Award (AY and JAS), Caroline Wiess Law Fund for Research in Molecular Medicine at Baylor College of Medicine (AY), National Hemophilia Foundation Innovative Investigator Research Award (AY), and Mary R. Gibson Foundation (AY). DG is a Howard Hughes Medical Institute Investigator.

## Supplemental Figure Legends

**Figure S1.**
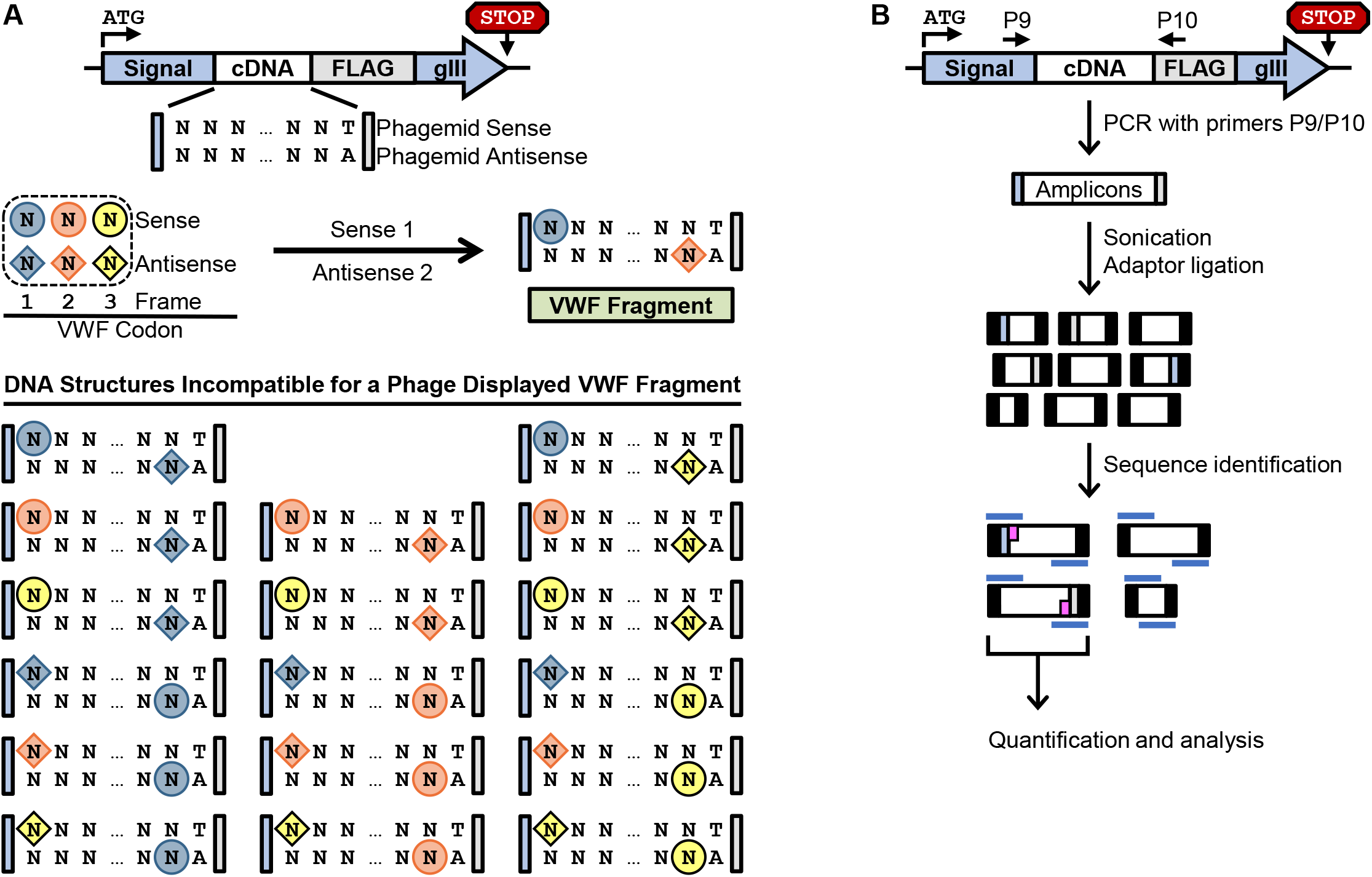
Schematic for expressing a VWF fragment fused to pIII and generating NGS-suitable DNA from the phage display library. A) Random fragments of the VWF plasmid (cDNA) were cloned into the phagemid. All possible combinations of orientation and frame are shown, but only the Sense 1/Antisense 2 combination can generate a displayed VWF fragment. This combination has an orientation in which the sense strand of the cloned VWF cDNA is in line with the sense strand of the phagemid and a frame that is bounded by the 1^st^ and 2^nd^ nucleotide of a VWF codon on the 5’ and 3’ ends, respectively. B) For NGS, PCR amplicons (~500bp-~1,200bp) generated with primers P9 and P10 risked inefficient and biased clustering onto the sequencing platform and were thus sonicated to ~100bp – ~600bp prior to ligation of Illumina-compatible, barcoded adaptors (black boxes). Paired end sequencing (blue bars, 75 or 100 bases) provided information on all adaptor-amplicon junctions. Only VWF fragment sequence (magenta) immediately adjacent to the vector sequence were analyzed.

**Figure S2.**
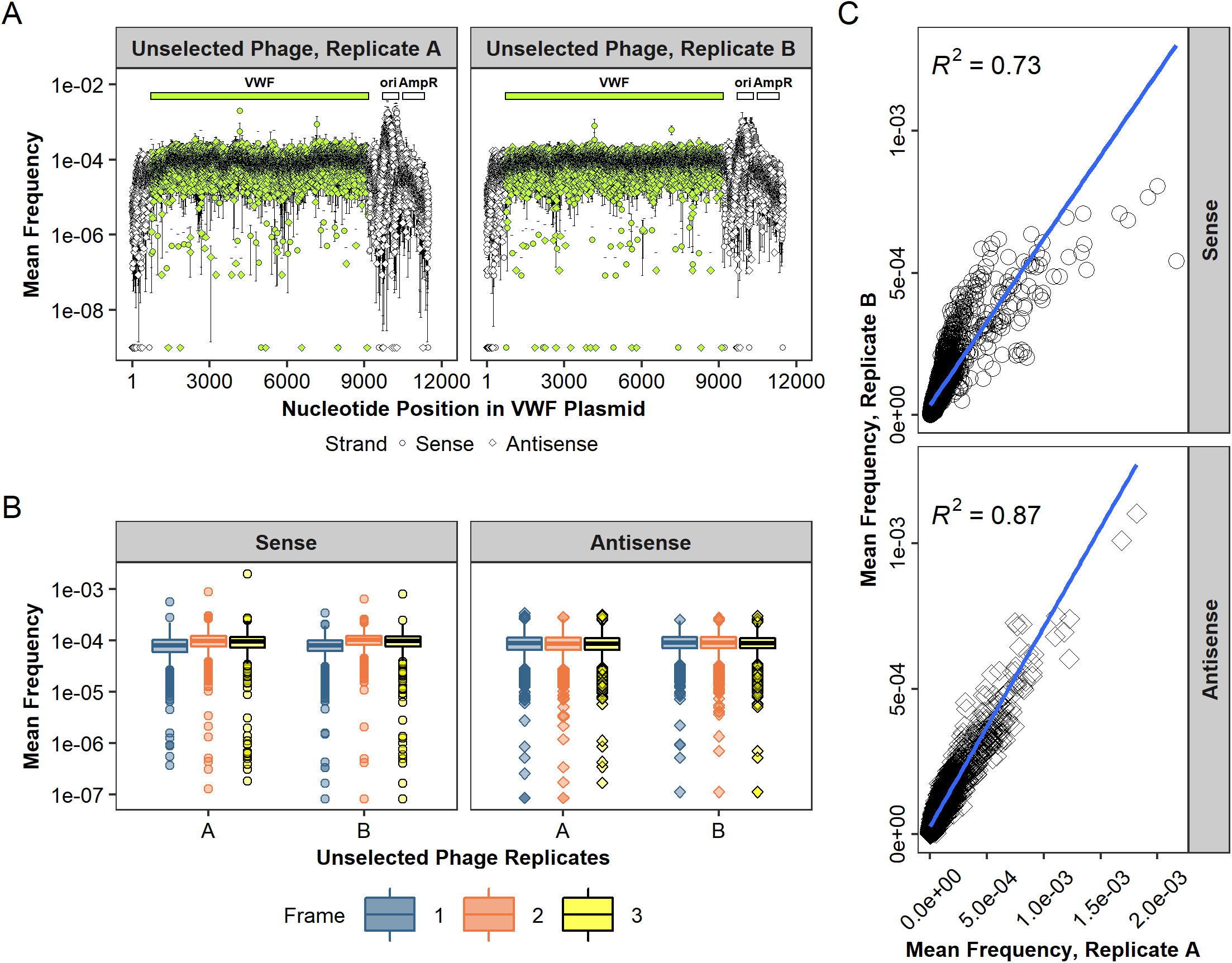
Packaging of VWF cDNA into phage virions is reproducible. A) The frequency (mean ± 1 standard deviation) of the nucleotides immediately adjacent to the phagemid adaptors (see Methods and Figure S1) was determined for the unselected phage (n = 3 NGS preparations for replicate A, n = 4 NGS preparations for replicate B) and mapped according to their position within the parental VWF plasmid. Each replicate of unselected phage was prepared from an aliquot of the expanded library stored at −80°C. Nucleotides with no reads are shown as a frequency of 1×10^−9^. Error bars represent 1 standard deviation. B) Boxplots of VWF nucleotides separated according to their strand (sense or antisense) and position within a VWF codon (Frame) show wide and differing distributions of frequencies but comparable medians. Nucleotides with no reads are not shown. C) The frequencies of each fragment terminus were well correlated between the unselected phage of the different selection experiments (Sense, Pearson R = 0.86; Antisense, Pearson R = 0.93).

## Notes

https://github.com/yeea00/anti-VWF_Phage.git

### Competing Interest Statement

The authors have declared no competing interest.

